# Age-dependent effects of intranasal oxytocin administration were revealed by resting brain entropy (BEN)

**DOI:** 10.1101/2024.11.25.625152

**Authors:** Donghui Song, Ze Wang

**Author notes:** Corresponding author: Donghui Song, No.19, Xinjiekouwai St, Haidian District, Beijing, 100875, China, Ze Wang, Ph.D., 670 W Baltimore St, Baltimore, MD, 21201, USA.

## Abstract

Brain entropy (BEN) reflects the irregularity, disorderliness, and complexity of brain activity and has gained increased interest in recent years. The demonstrated sensitivity of BEN to caffeine and medicine suggests the existence of neurochemical effects of BEN. Oxytocin (OT), a neuropeptide associated with childbirth and lactation, affects both social behavior and brain activity. The purpose of this study was to examine whether OT affects BEN in young and old adults.

A randomized, double-blind, placebo-controlled, two-factor (Age × OT) between-subjects design was used, and a total of seventy-five eligible healthy participants were included in the experiment. In the young adult group (YA), 23 participants received intranasal OT administration, while 18 received a placebo (PL) administration. In the older adult group (OA), 16 participants received intranasal OT administration, and 18 received PL administration.

Using fMRI-based BEN mapping, we found the age-dependent effect of intranasal OT in the left temporal parietal junction (TPJ), where BEN increased in YA and BEN decreased in OA. A whole-brain functional connectivity (FC) analysis with the left TPJ as the seed and we found that FC between the left TPJ and right TPJ increased in both YA and OA. FC of left and right TPJ and plasma OT contribute to left TPJ BEN just found in YA with intranasal OT administration. These results indicate that BEN is sensitive to age-related effects of neurochemical signals and highlight plasma OT on the effects of intranasal OT in young adults.

## 1. Introduction

Brain entropy (BEN) reflects the irregularity, disorderliness and complexity of brain activity (Wang, Li et al. 2014), garnering increasing attention due to its ability to capture patterns relevant to brain function and diseases across different data modalities and time scales (Stam 2005, Wang, Li et al. 2014, Wang and Initiative 2020, Song, Jann et al. 2024). Resting-state magnetic resonance imaging (rs-fMRI)-based BEN offers unique advantages in localizing brain function compared to electroencephalography (EEG) and magnetoencephalography (MEG), owing to its superior spatial resolution. Compared to other rs-fMRI metrics such as the fractional amplitude of low-frequency fluctuations (fALFF) (Zou, Zhu et al. 2008) and cerebral blood flow (CBF) (Li, Zhu et al. 2012), BEN offers complementary information for detecting brain activity (Song, Chang et al. 2019). Importantly, the relationship between BEN, cognition, and task activation has been established (Wang 2021, Lin, Chang et al. 2022, Del Mauro and Wang 2024, Song and Wang 2024). Aberrant BEN patterns have been identified across various brain disorders, including depression (Lin, Lee et al. 2019, Liu, Song et al. 2020, Dong-Hui Song 2024), addiction disorders (Li, Fang et al. 2016, Chang, Zhang et al. 2018, Jiang, Cai et al. 2023), Alzheimer’s disease (Wang and Initiative 2020), schizophrenia (Xue, Yu et al. 2019) et al. Furthermore, BEN demonstrates sensitivity to neuroplasticity, including responses to non-invasive brain stimulation (Song, Chang et al. 2019, Liu, Song et al. 2024) and caffeine (Chang, Song et al. 2018). Recently, we found that BEN is associated with ovarian hormones (progesterone) and BEN mediated the relationship between progesterone and behavioral activation systems (Song and Wang 2024). These findings suggest that BEN can effectively capture the impact of neurochemical molecules on brain function. However, research on how neurochemical molecules affect BEN remains limited. This study will further explore the effects of neurochemical molecules on BEN by administering oxytocin (OT) via intranasal inhalation. OT is a peptide hormone and neuropeptide, which plays a key role in the reproductive system, including childbirth and lactation (Lee, Macbeth et al. 2009). Over the past decade, due to its influence on human social behaviors, it has garnered widespread attention (Neumann 2008, Bartz, Zaki et al. 2011, Froemke and Young 2021). OT is primarily synthesized in the hypothalamus, it then projects to other brain regions as a neuromodulator impacting various social behaviors, including social perception and memory, affiliation, aggression, and empathy (Neumann 2008, Bartz, Zaki et al. 2011, Froemke and Young 2021). However, the effects of OT on the brain remain puzzling (Grace, Rossell et al. 2018, Quintana, Lischke et al. 2021). Recent studies indicate that OT can cause age-dependent effects on brain function from the task and rs-fMRI (Horta, Pehlivanoglu et al. 2020). Campbell et al. (2014) found older males receiving OT showed improved emotion recognition relative to those in the placebo (PL) group and no differences were found for older females or young adults (Campbell, Ruffman et al. 2014). Horta et al. (2019) found that OT altered the processing of sadness and happiness for young participants, while OT only affected the processing of sadness for older participants (Horta, Ziaei et al. 2019). Frazier et al. (2021) found that OT doesn’t modulate the behavioral effects in their investment after breach-of-trust feedback between younger and older participants. However, after breach-of-trust feedback, older participants in the OT group showed less activity in the left superior temporal gyrus than in the PL group (Frazier, Lin et al. 2021). Based on rs-fMRI, Liu et al. (2022) found that intranasal OT administration reduced functional connectivity (FC) between the amygdala and the ventral salience network for older but not young adults (Liu, Lin et al. 2022). Recently, Xiao et al. (2024) found that intranasal OT decreased the connectivity density and strength in the thalamus for young participants, while intranasal OT increased the connectivity density and strength in the caudate for older participants. These studies indicate that OT causes age-related effects on brain function. Del Mauro et al. (2024) recently found that pain intensity has different BEN patterns across age groups. Specifically, pain intensity was negatively correlated with BEN in the somatosensory and occipital cortex in young adults. In contrast, in the middle-aged and aging group, pain intensity showed a positive association with increased BEN in the somatomotor cortex, cingulate cortex, precuneus, inferior parietal lobules, prefrontal cortex, anterior insula, occipital cortex, cerebellum (Del Mauro, Sevel et al. 2024). Existing literature has shown that OT can effectively alleviate pain sensitivity (Rash, Aguirre-Camacho et al. 2014, Tracy, Georgiou-Karistianis et al. 2015, Zunhammer, Geis et al. 2015, Herpertz, Schmitgen et al. 2019). These studies combined with the growing literature that BEN can capture age-related changes in brain function (Del Mauro, Sevel et al. 2024, Zhao, Shuai et al. 2024), suggest that BEN is highly likely to capture the age-dependent effects caused by OT.

In conclusion, BEN is available to capture age-dependent differences induced by OT and we aim to address this question using publicly available data from OpenNeuro ds004725 (Markiewicz, Gorgolewski et al. 2021).

## 2. Methods

### 2.1 Participants

The dataset from OpenNeuro ds004725 (https://openneuro.org/datasets/ds004725/versions/1.0.1) was collected between August 2013 and October 2014. Total of eighty-seven, white, English-speaking generally healthy adults, including 44 healthy young adults (YA, aged 18-31 years; 21 females) and 43 older adults (OA, aged 63-81 years; 24 females) without neurological or psychiatric disorders, and all participants provided written informed consent for the original study (Liu, Lin et al. 2022).

### 2.2 Experimental Procedure

A randomized, double-blind, placebo-controlled, between-subject design followed standardized procedures for intranasal oxytocin self-administration (Guastella, Hickie et al. 2013). The PL contained identical ingredients except for the OT. All study personnel involved in recruitment and scheduling and who engaged directly with study participants, and the participants themselves, were blinded to the treatment conditions.

The participants aged 18-31 and 63-81 were prescreened via phone. Eligible participants then came to the University of Florida (UF) for an in-person screening session, during the in-person screening session, participants completed an intake interview and cognitive measures that included the Digit Symbol Substitution Test (DSST), which measures sensorimotor processing speed (Wechsler 1981), and the Rey Auditory Verbal Learning task (RAVLT), which measures short-term verbal memory (Rey 1958). Plasma OT was also collected by blood sampling (Horta, Polk et al. 2023). Eligible participants self-administer a single dose (24 IUs) to administer the targeted dose of 24 IUs of either OT or PL via nasal spray. Promptly after OT or PL administration, participants were transported to the imaging facility and settled into the MRI scanner. T1-weighted anatomical images were acquired and the rs-fMRI scan. During the resting-state scan, which took place 70–90 min after spray administration, participants were asked to relax and look at a white fixation cross on a black screen. All participants were instructed to avoid food, exercise, and sexual activity for 2 h and smoking, caffeine, alcohol, and recreational drugs for 24 hours before the visit.

### 2.3 MRI acquisition

The MRI data were acquired on a 3T Philips Achieva MR Scanner (Philips Medical Systems, Best, The Netherlands) with a 32-channel head coil. A high-resolution structural image was acquired using a magnetization-prepared rapid gradient echo (MP-RAGE) sequence with parameters: field of view (FOV) = 240 mm x 240 mm x 240 mm, in-plane resolution = 1 mm × 1 mm, slice thickness = 1 mm without gap, sagittal slice orientation. The rs-fMRI images were acquired using a gradient-echo-planar imaging (EPI) sequence with parameters: repetition time (TR) = 2 s, echo time (TE) = 30 ms, FOV = 252 mm x 252 mm x 133 mm, flip angle = 90°, transverse slice orientation, in-plane resolution = 3.15×3.15 mm, slice thickness = 3.5 mm without skip with a total of 38 interleaved slices. A total of 240 volumes were acquired during the 8-minute.

For more detailed participants’ information, the experimental procedure, and MRI acquisition parameters can be found in the original article from the dataset (Liu, Lin et al. 2022, Horta, Polk et al. 2023).

### 2.4 MRI preprocessing

MRI data were preprocessed using the same configuration and pipeline as in our previous study (Liu, Song et al. 2024, Song, Deng et al. 2024, Song and Wang 2024). The fMRIPrep (Esteban, Markiewicz et al. 2019) (version=23.1.4) based on Nipype (version= 1.8.6) (Gorgolewski, Burns et al. 2011), that containerized to docker (version=24.0.7) (https://www.docker.com/) (https://pypi.org/project/fmriprep-docker/) was used to perform pre processing under Ubuntu (version=22.04.3 LTS) (https://releases.ubuntu.com/jammy/). De noising was performed using customed python (version=3.10) (https://www.python.org/) s cripts that are based-Nilearn (version=0.10.2) (https://nilearn.github.io/stable/index.html).

The structural images were skull-stripped and segmented into gray matter (GM), white matter (WM), and cerebrospinal fluid (CSF) using FAST (Zhang, Brady et al. 20 01), then normalized to MNI space. The functional images underwent preprocessing through the following steps: (1) First, a reference volume and its skull-stripped version were generated using a custom methodology of fMRIPrep. (2) Head motion correction was performed using mcflirt (Jenkinson, Bannister et al. 2002). (3) The functional ref erence was then co-registered to the T1w reference using mri_coreg followed by flirt (Jenkinson and Smith 2001) and framewise displacement (FD) (Power, Barnes et al. 2012), and DVARS were calculated. The average signals were extracted within the GM, CSF, and WM. (4) The functional images were resampled into MNI space using antsApplyTransforms (ANTs). (5) Detrending and temporal bandpass filtering (0.009–0.08 Hz) was performed. (6) Nuisance components were regressed, including the six motion parameters, FD, standardized DVARS, root mean square displacement, and the average WM signals and average CSF signals. (7) Smoothing with an isotropic Gaussian kernel (FW HM□ = □6 mm).

### 2.5 Gray matter volume

It has been widely acknowledged that there is gray matter volume (GMV) decreased in elderly individuals (Good, Johnsrude et al. 2001, Hafkemeijer, Altmann-Schneider et al. 2014, Bethlehem, Seidlitz et al. 2022). Furthermore, recent research found a brain-wise negative correlation between GMV and BEN (Del Mauro and Wang 2024). To mitigate the potential impact of GMV on BEN, we computed GMV from structural images of each participant. GMV was computed by the fMRIPrep-FSL pipeline, the pipeline has been validated to be equally effective with the Computational Anatomy Toolbox (CAT) 12.8 pipeline (Gaser, Dahnke et al. 2022) in reflecting age information (Antonopoulos, More et al. 2023). Firstly, the GM was segmented from the fMRIPrep pipeline (see 2.4), then following FSLVBM protocol (https://fsl.fmrib.ox.ac.uk/fsl/fslwiki/FSLVBM), fslvbm_2_template was used to create the study-specific GM template, fslvbm_3_proc was used to register all GM images to the study-specific template non-linearly, then the GM images were smoothed using an isotropic Gaussian kernel with FWHM□=□4 mm. The detailed guide of FSLVBM is available at https://fsl.fmrib.ox.ac.uk/fsl/fslwiki/FSLVBM/UserGuide. The total GMV of the whole brain was calculated using a script from http://www0.cs.ucl.ac.uk/staff/gridgway/vbm/get_totals.m.

### 2.6 BEN mapping

The voxel-wise BEN maps were calculated from preprocessed rs-fMRI data by the BEN mapping toolbox (BENtbx) (Wang, Li et al. 2014) based on sample entropy (Richman and Moorman 2000). The toolbox is available at https://www.cfn.upenn.edu/zewang/BENtbx.php and https://github.com/zewangnew/BENtbx. The window length was set to 3 and the cutoff threshold was set to 0.6 according to our experimental optimization (Wang, Li et al. 2014). The first four volumes were discarded for signal stability, BEN maps were smoothed with an isotropic Gaussian kernel with FWHM□=□8 mm. More details of BEN calculation can be found in our previous studies (Wang, Li et al. 2014, Song, Chang et al. 2019, Song, Chang et al. 2019, Wang 2021, Lin, Chang et al. 2022).

### 2.7 Functional connectivity

We selected the brain regions that showed significant BEN effects as regions of interest (ROI) to assess the distance effects of OT to perform seed-based FC analysis. The seed-based FC calculation was as follows: The ROI was binarized to create a mask and utilized as a seed. Using preprocessed rs-fMRI images, FC maps were calculated using custom Python scripts based on Nilearn (https://nilearn.github.io/stable/index.html). The mean value of the rs-fMRI time series was extracted from the seed and correlated to the time series of all voxels in the whole brain. Pearson’s correlation coefficient was measured as the amplitude of FC and Fisher’s z-transformed to improve the normality before performing group-level analysis. This analytical process can also be seen in our previous studies (Song, Chang et al. 2019, Dong-Hui Song 2024, Song and Wang 2024).

## 3. Results

### 3.1 Quality control and information of participants

Twelve participants were removed from the dataset, two of whom had shorter volumes (190 volumes), one without values of plasma OT, and nine of whom had large head motion (mean FD > 0.5 mm). Finally, seven-five participants were included in the study, one of the participants is left-handed, while the others are all right-handed, the YA group included 23 OT (age=21.81±2.72 years,11 females) and 18 PL participants (age=23.37±3.16 years,9 females), the OA group included 16 OT (age=71.44±5.58 years,10 females), and PL included 18 (age=71.78±4.48 years,11females).

Two-way analysis of variance (ANOVA) was used to detect group differences in age (YA and OA) and treatment (OT and PL), as well as their interaction effects for education (Edu), body mass index (BMI), DSST, RAVLT, plasma OT, Mean FD, and Total GMV. The results indicated significant differences between the YA and OA groups in Edu, DSST, RAVLT and Mean FD. Significant differences were also found between OT and PL in BMI. No significant interaction effects were observed between age and treatment (see Table 1).

**Table 1.**
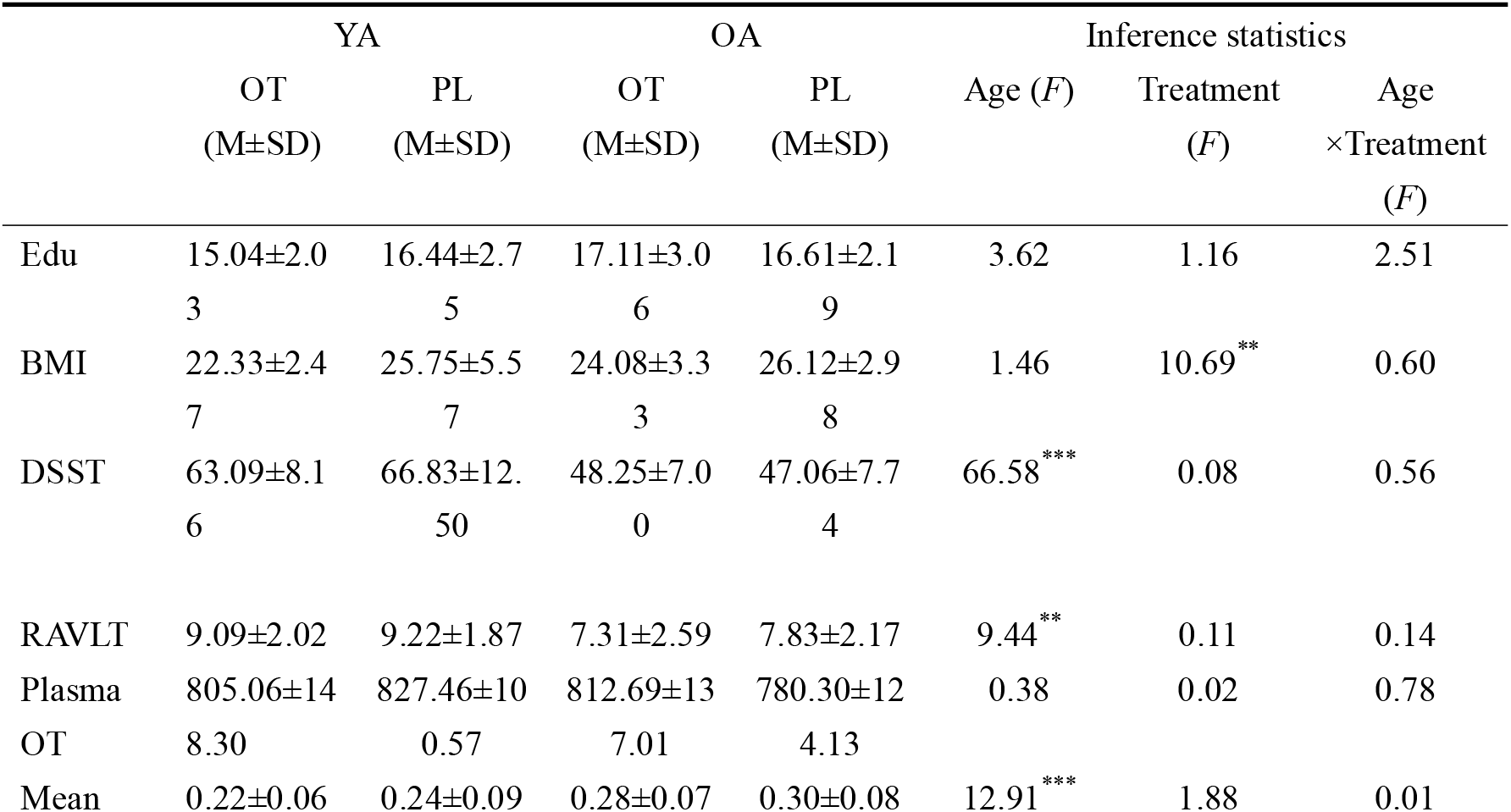
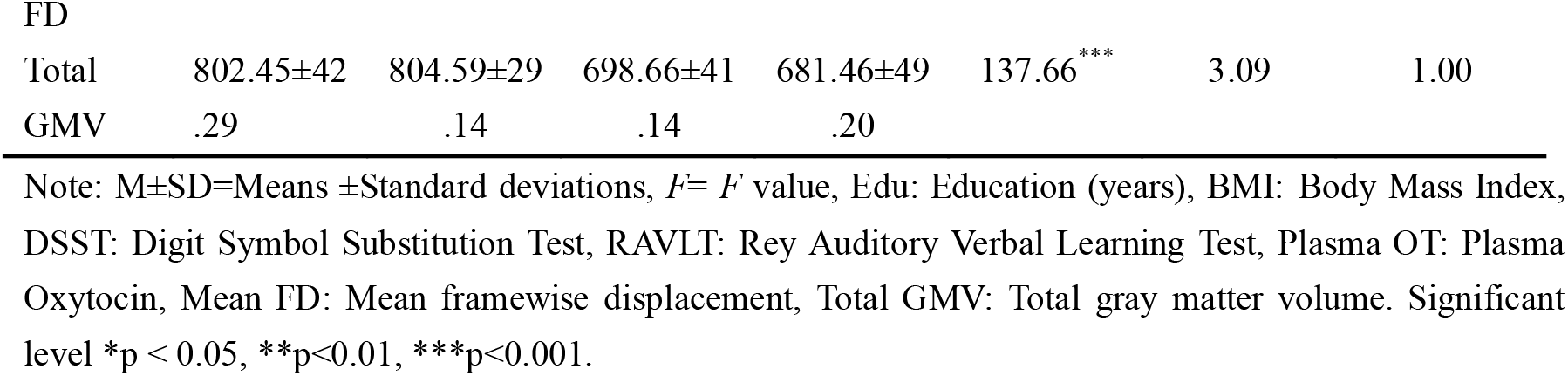
Participants’ information and inference statistics (age, treatment, and age × treatment effects) for education, BMI, cognitive, Plasma OT, Mean FD, and Total GMV (N = 75)

### 3.2 Age-dependent effects of intranasal oxytocin administration in left TPJ

Statistical Parametric Mapping (SPM12, WELLCOME TRUST CENTRE FOR NEUROIMAGING, London, UK, http://www.fil.ion.ucl.ac.uk/spm/software/spm12/) (Friston, Holmes et al. 1994) was used to perform voxel-wise statistical analysis. We employed a two-way analysis of covariance (ANCOVA) to assess the effects between age and treatment, with age, gender, handedness, Edu, BMI, DSST, RAVLT, Mean FD and Total GMV included as covariates. The results showed a significant interaction between age and treatment on BEN in the left TPJ (*F* _(1,62)_>11.93, voxel-wise *p* < 0.001, cluster size > 20). Subsequently, we extracted the mean BEN of the left TPJ from each participant. Two sample t-tests were then performed on YA and OA groups respectively after regressing out age, gender, handedness, Edu, BMI, DSST, RAVLT, Mean FD, and Total GMV. The results indicated that in the YA group, OT increased BEN in the left TPJ relative to PL (*t*_*(23,18)*_=3.05, *p*=0.004), whereas, in the OA group, OT decreased BEN in the left TPJ relative to PL (*t*_*(16,18)*_ =-3.84, *p*<0.001), we also found that BEN in the OA group are higher than those in the YA group for participants who received PL administration (*t*_*(18,18)*_=3.80, *p*<0.001), while the BEN in the YA group is higher than those in the OA group for participants who received OT administration (*t*_*(23,16)*_=3.04, *p*=0.004) (see Fig1. A).

**Fig 1.**
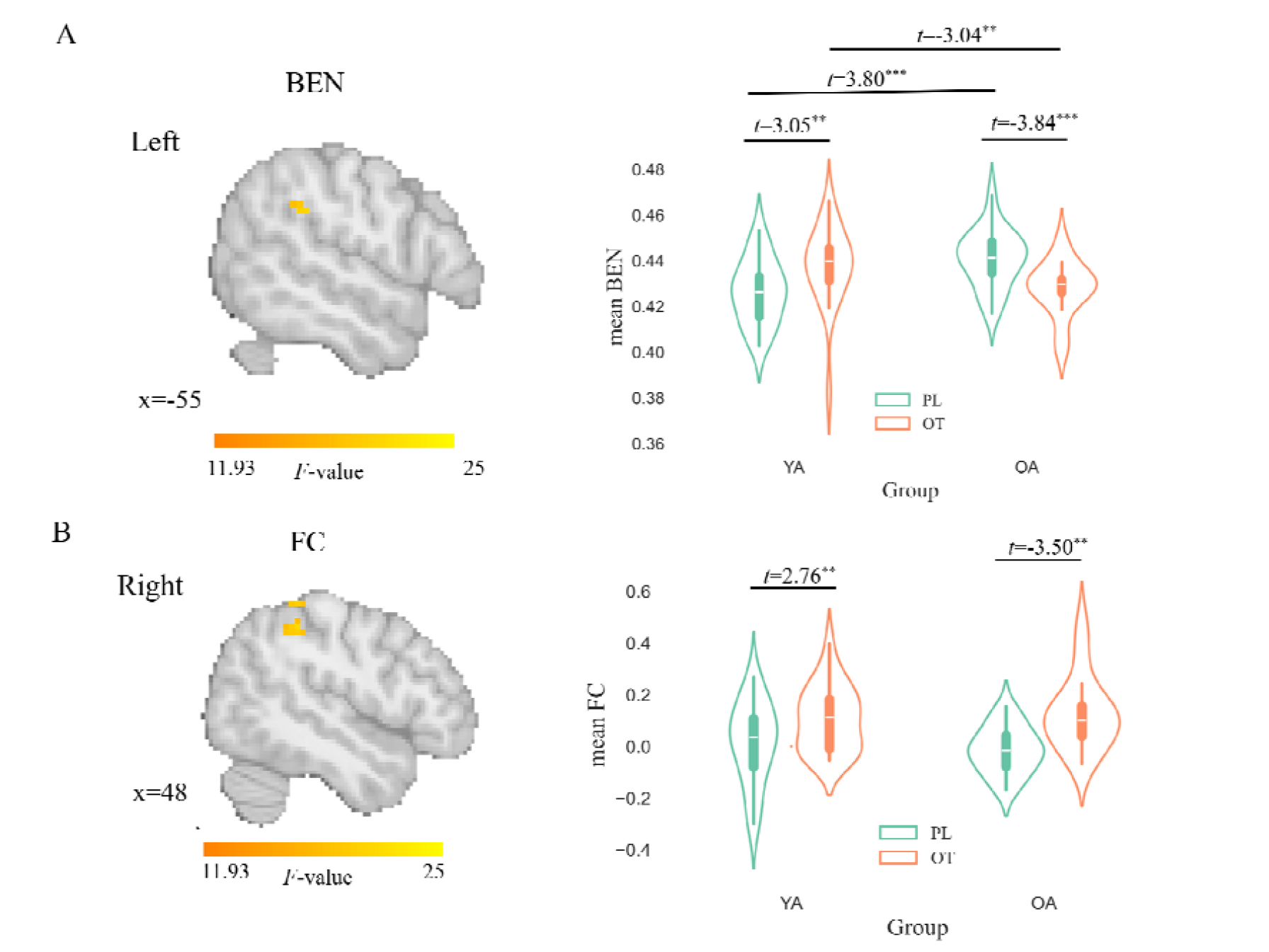
Age-dependent effects of intranasal OT administration in left TPJ. A: The interaction effects of age (YA vs OA) and treatment (OT vs PL) on BEN in left TPJ. Left showed the brain region of interaction effects. Left indicates the left hemisphere. Colorbar indicates *F* value. The number indicates location along the x-axis in the MNI space of brain slice. Right showed that mean BEN of left TPJ for each subgroup and the significance of the difference. The t indicates value after regressing out covariances. The y-axis represents the mean BEN from the left TPJ. B: The main effects of treatment (OT vs PL) on left TPJ FC in right TPJ. Left showed the brain region of main effects. Right indicates the right hemisphere. Colorbar indicates *F* value. The number indicates location along the x-axis in the MNI space of brain slice. Right showed that mean FC of right TPJ for each subgroup and the significance of the difference. The t indicates value after regressing out covariances. The y-axis represents the mean FC from the right TPJ. Significance level * *p*<0.05, ** *p*<0.01, ** *p*<0.001.

We also analyzed whole-brain FC seeded from the left TPJ using ANCOVA. The results revealed a significant main effect of treatment in the right TPJ (*F* _(1,62)_>11.93, voxel-wise *p*<0.001, FWE corrected *p*<0.05) and significant increases in FC between the left TPJ and right TPJ in both the YA (*t*_*(23,18)*_=2.76, *p*=0.009) and OA groups (*t*_*(16,18)*_ =3.50, *p*=0.001) (see Fig1. B). Table 2 shows cluster information for ANCOVA.

**Table 2.** Clusters size table for ANCOVA.

| Contrast | Cluster ID | Peak MNI<br>Coordinate (X, Y,<br>Z) | Peak<br>Value (*F*) | Cluster Size | Brain Region |
| --- | --- | --- | --- | --- | --- |
| BEN ANCOVA | 1 | (-60, -39,30) | 20.79 | 28 | L TPJ |
| FC ANCOVA | 1a | (45, -42,63) | 18.07 | 34 | R TPJ |
|  | 1b | (48, -39,48) | 17.22 |  |  |
|  | 1c | (39, -45,69) | 14.19 |  |  |

### 3.3 Plasma OT and FC contribute to BEN in left TPJ in YA with intranasal oxytocin administration

To examine the relationship between plasma OT and FC of left-right TPJ with left TPJ BEN, we performed linear regression analyses for each subgroup, using the residuals of plasma OT (covariates including age, gender, handedness, Edu, BMI, DSST, RAVLT, and Total GMV), left-right TPJ FC (covariates including age, gender, handedness, Edu, BMI, DSST, RAVLT, Mean FD, and Total GMV), and left TPJ BEN (covariates including age, gender, handedness, Edu, BMI, DSST, RAVLT, Mean FD, and Total GMV) after regressing out the covariates.

The results show that FC of the left-right TPJ (*t*_*23*_ = 2.25, *p* = 0.036) and plasma OT (*t*_*23*_ = -2.82, *p* = 0.011) significantly predicted left TPJ BEN in the YA group with intranasal OT administration (see Table 3). The regression model’s adjusted *R*^*2*^ = 0.37 (*F* _(2, 20)_ = 7.45, *p* = 0.004) was. However, no predictive effects of FC of left-right TPJ and plasma OT on left TPJ BEN were found in the other groups.

**Table 3.** FC and plasma OT contribute to BEN in left TPJ in YA with OT administration.

| variables | $\beta$ | $t$ | $p$ | CI [0.025,0.975] |
| --- | --- | --- | --- | --- |
| FC | 52.8655 | 2.25 | 0.036 | 3.900, 101.831 |
| Plasma OT | -0.0551 | -2.82 | 0.011 | -0.096, -0.014 |
| Intercept | 2.2625 | 0.81 | 0.427 | 3.563, 8.088 |

Note : CI: confidence interval

## 4. Discussion

The results indicated that BEN detects the age-dependent effects of intranasal OT. Specifically, a decreased BEN in the left TPJ in the OA group, while BEN increases in the left TPJ in the YA group. Further, seed-based FC analysis using the left TPJ as a seed revealed increased FC between the left and right TPJ in both YA and OA groups. Interestingly, plasma OT and FC of left-right TPJ showed significant prediction for BEN in the left TPJ just in the YA group with intranasal OT and no such effect was observed in the OA group. The coupling of plasma OT, FC of left-right TPJ and BEN in the left TPJ in the YA group and its decoupling in the OA group suggest the possible reason for the age-dependent effects of intranasal OT.

The TPJ plays a crucial role in various cognitive functions, primarily including attention (Young, Dodell-Feder et al. 2010, Geng and Vossel 2013), social cognition (Van Overwalle 2009, Krall, Rottschy et al. 2015), and multisensory integration (Ionta, Heydrich et al. 2011, Kheradmand and Winnick 2017), serving as a core hub in the social brain network (Blakemore 2008, Van Overwalle 2009, Wang, Metoki et al. 2021). Intranasal OT has garnered extensive attention for its effects on social attention and cognition and numerous studies have demonstrated its facilitative role in encoding, consolidation, and recognition of social information across various contexts (Bartz, Zaki et al. 2011, Shahrestani, Kemp et al. 2013, Horta, Pehlivanoglu et al. 2020, Quintana, Lischke et al. 2021). Existing literature suggests that the effects of intranasal OT are closely related to the TPJ (Rocchetti, Radua et al. 2014, Hu, Scheele et al. 2016, Grace, Rossell et al. 2018).

In the study, intranasal OT led to decreased BEN in the left TPJ and FC between the left TPJ and the right TPJ was increased in the OA group. It is noteworthy that in the PL, OA exhibit higher BEN in the left TPJ compared to YA. Our previous studies indicated increased BEN in normal aging (Wang and Initiative 2020, Del Mauro, Sevel et al. 2024) suggesting that decreasing BEN in the left TPJ is beneficial for older adults. Intranasal OT has been shown to alleviate pain (Bos, Montoya et al. 2015, Boll, De Minas et al. 2018, Lussier, Cruz-Almeida et al. 2019, Kawasaki, Sakai et al. 2024). Recently our study found that higher pain intensity is positively correlated with BEN in TPJ in middle-aged and older adults (Del Mauro, Sevel et al. 2024). This suggests that decreased BEN in left TPJ by intranasal OT attenuates pain in elderly individuals. We also observed increased FC of left TPJ with right TPJ in the OA group. Given that older adults exhibit reduced attentional efficiency and diminished inhibitory control of attention (Commodari and Guarnera 2008), this increased FC may suggest enhanced communication between external perception and awareness, leading to improved selective attention and sustained control capabilities (Ebner, Horta et al. 2015).

It is intriguing that among the YA group, intranasal OT elevated BEN in the left TPJ, whereas increased FC between the left TPJ and right TPJ. We also found that BEN in the left TPJ was predicted by plasma OT and FC of left-right TPJ in the YA with intranasal OT, but in the OA. This could be a key factor contributing to the age-dependent effects of intranasal OT. Lancaster et al. (2015) found that plasma OT robustly predicted activation in areas critical for social cognitive processes including TPJ, and higher levels of plasma OT are associated with increased activity in TPJ which are important for social cognitive processes in YA (Lancaster, Carter et al. 2015). Although it is difficult for plasma OT to cross the blood-brain barrier, studies have shown that intranasal OT increases both plasma (peripheral) and central OT levels. The peripheral and central OT are positive correlations particularly after intranasal OT, with the correlation reaching as high as 0.66 (Valstad, Alvares et al. 2017). Furthermore, research indicates that the effects of intranasal OT are dose-dependent (Cardoso, Ellenbogen et al. 2013, Leng and Ludwig 2016, Quintana, Westlye et al. 2017, Martins, Brodmann et al. 2022), with different plasma OT interacting with varying doses of intranasal OT in YA.

Several limitations need to be addressed. First, the experimental design is between-subjects, which means there may be substantial variability between participants, potentially affecting the results. Secondly, the sample size is relatively small for the effects of plasma OT, and the timing of plasma OT sampling varied from a few days to several weeks before the scan. Although the study suggests that baseline plasma OT remains relatively stable over 6–9 months (Feldman, Weller et al. 2007), intranasal OT can cause a significant increase in plasma OT (Valstad, Alvares et al. 2017). Therefore, the impact of baseline plasma OT on the effects of intranasal OT administration warrants further investigation in larger samples. In summary, the study reveals that BEN is sensitive to age-related neurochemical influences and underscores the impact of plasma OT on the effects of intranasal OT in young adults.

## Acknowledgments

We thank Marilyn Horta, Rebecca Polk, and Natalie Ebner et al for sharing their data.

## Data and code availability

The original data is available in the online repository of OpenNeuro ds004725 (https://openneuro.org/datasets/ds004725/versions/1.0.1). The GMV maps, BEN maps, FC maps, and unthresholded statistical maps can be accessed from OSF (https://osf.io/6d73g/?view_only=e3f279fa1ff34c58a07b560a9746dff2) to repeat the study. The fmriprep is available at https://fmriprep.org/en/0.6.4/index.html. Nilearn is available at https://nilearn.github.io/dev/index.html. FSL is available at https://fsl.fmrib.ox.ac.uk/fsl/docs/#/. The total GMV calculation is available at http://www0.cs.ucl.ac.uk/staff/gridgway/vbm/get_totals.m. SPM is available at https://www.fil.ion.ucl.ac.uk/spm/software/spm12/. BENtbx is available at https://www.cfn.upenn.edu/zewang/BENtbx.php and https://github.com/zewangnew/BENtbx.

## CRediT authorship contribution statement

Donghui Song: conceptualization, data analysis, visualization, manuscript drafting, and editing. Ze Wang: conceptualization, manuscript review and editing, supervision.

## Notes

https://openneuro.org/datasets/ds004725/versions/1.0.1

### Competing Interest Statement

The authors have declared no competing interest.

## References

Antonopoulos, G., S. More, F. Raimondo, S. B. Eickhoff, F. Hoffstaedter and K. R. Patil (2023). “A systematic comparison of VBM pipelines and their application to age prediction.” Neuroimage 279: 120292.

Bartz, J. A., J. Zaki, N. Bolger and K. N. Ochsner (2011). “Social effects of oxytocin in humans: context and person matter.” Trends in cognitive sciences 15(7): 301–309.

Bethlehem, R. A., J. Seidlitz, S. R. White, J. W. Vogel, K. M. Anderson, C. Adamson, S. Adler, G. S. Alexopoulos, E. Anagnostou and A. Areces-Gonzalez (2022). “Brain charts for the human lifespan.” Nature 604(7906): 525–533.

Blakemore, S.-J. (2008). “The social brain in adolescence.” Nature Reviews Neuroscience 9(4): 267–277.

Boll, S., A. A. De Minas, A. Raftogianni, S. Herpertz and V. Grinevich (2018). “Oxytocin and pain perception: from animal models to human research.” Neuroscience 387: 149–161.

Bos, P. A., E. R. Montoya, E. J. Hermans, C. Keysers and J. van Honk (2015). “Oxytocin reduces neural activity in the pain circuitry when seeing pain in others.” Neuroimage 113: 217–224.

Campbell, A., T. Ruffman, J. E. Murray and P. Glue (2014). “Oxytocin improves emotion recognition for older males.” Neurobiology of aging 35(10): 2246–2248.

Cardoso, C., M. A. Ellenbogen, M. A. Orlando, S. L. Bacon and R. Joober (2013). “Intranasal oxytocin attenuates the cortisol response to physical stress: a dose–response study.” Psychoneuroendocrinology 38(3): 399–407.

Chang, D., D. Song, J. Zhang, Y. Shang, Q. Ge and Z. Wang (2018). “Caffeine caused a widespread increase of resting brain entropy.” Scientific reports 8(1): 2700.

Chang, D., J. Zhang, W. Peng, Z. Shen, X. Gao, Y. Du, Q. Ge, D. Song, Y. Shang and Z. Wang (2018). “Smoking cessation with 20 Hz repetitive transcranial magnetic stimulation (rTMS) applied to two brain regions: a pilot study.” Frontiers in human neuroscience 12: 344.

Commodari, E. and M. Guarnera (2008). “Attention and aging.” Aging clinical and experimental research 20: 578–584.

Del Mauro, G., L. S. Sevel, J. Boissoneault and Z. Wang (2024). “Divergent association between pain intensity and restingLstate fMRILbased brain entropy in different age groups.” Journal of Neuroscience Research 102(5): e25341.

Del Mauro, G. and Z. Wang (2024). “Associations of brain entropy estimated by resting state fMRI with physiological indices, body mass index, and cognition.” Journal of Magnetic Resonance Imaging 59(5): 1697–1707.

Del Mauro, G. and Z. Wang (2024). “rsfMRI-based Brain Entropy is negatively correlated with Gray Matter Volume and Surface Area.” bioRxiv: 2024.2004. 2028.591371.

Dong-Hui Song, Y.W., Ze Wang (2024). “Increased Resting Brain Entropy in Mild to Moderate Depression was Decreased by Nonpharmacological Treatment “ submitted.

Ebner, N. C., M. Horta, T. Lin, D. Feifel, H. Fischer and R. A. Cohen (2015). “Oxytocin modulates meta-mood as a function of age and sex.” Frontiers in Aging Neuroscience 7: 175.

Esteban, O., C. J. Markiewicz, R. W. Blair, C. A. Moodie, A. I. Isik, A. Erramuzpe, J. D. Kent, M. Goncalves, E. DuPre and M. Snyder (2019). “fMRIPrep: a robust preprocessing pipeline for functional MRI.” Nature methods 16(1): 111–116.

Feldman, R., A. Weller, O. Zagoory-Sharon and A. Levine (2007). “Evidence for a neuroendocrinological foundation of human affiliation: plasma oxytocin levels across pregnancy and the postpartum period predict mother-infant bonding.” Psychological science 18(11): 965–970.

Frazier, I., T. Lin, P. Liu, S. Skarsten, D. Feifel and N. C. Ebner (2021). “Age and intranasal oxytocin effects on trust-related decisions after breach of trust: Behavioral and brain evidence.” Psychology and Aging 36(1): 10.

Friston, K. J., A. P. Holmes, K. J. Worsley, J. P. Poline, C. D. Frith and R. S. Frackowiak (1994). “Statistical parametric maps in functional imaging: a general linear approach.” Human brain mapping 2(4): 189–210.

Froemke, R. C. and L. J. Young (2021). “Oxytocin, neural plasticity, and social behavior.” Annual Review of Neuroscience 44: 359–381.

Gaser, C., R. Dahnke, P. M. Thompson, F. Kurth, E. Luders and A. s. D. N. Initiative (2022). “CAT–A computational anatomy toolbox for the analysis of structural MRI data.” biorxiv: 2022.2006. 2011.495736.

Geng, J. J. and S. Vossel (2013). “Re-evaluating the role of TPJ in attentional control: contextual updating?” Neuroscience & Biobehavioral Reviews 37(10): 2608–2620.

Good, C. D., I. S. Johnsrude, J. Ashburner, R. N. Henson, K. J. Friston and R. S. Frackowiak (2001). “A voxel-based morphometric study of ageing in 465 normal adult human brains.” Neuroimage 14(1): 21–36.

Gorgolewski, K., C. D. Burns, C. Madison, D. Clark, Y. O. Halchenko, M. L. Waskom and S. S. Ghosh (2011). “Nipype: a flexible, lightweight and extensible neuroimaging data processing framework in python.” Frontiers in neuroinformatics 5: 13.

Grace, S. A., S. L. Rossell, M. Heinrichs, C. Kordsachia and I. Labuschagne (2018). “Oxytocin and brain activity in humans: a systematic review and coordinate-based meta-analysis of functional MRI studies.” Psychoneuroendocrinology 96: 6–24.

Guastella, A. J., I. B. Hickie, M. M. McGuinness, M. Otis, E. A. Woods, H. M. Disinger, H.-K. Chan, T. F. Chen and R. B. Banati (2013). “Recommendations for the standardisation of oxytocin nasal administration and guidelines for its reporting in human research.” Psychoneuroendocrinology 38(5): 612–625.

Hafkemeijer, A., I. AltmannLSchneider, A. J. de Craen, P. E. Slagboom, J. van der Grond and S. A. Rombouts (2014). “Associations between age and gray matter volume in anatomical brain networks in middleLaged to older adults.” Aging cell 13(6): 1068–1074.

Herpertz, S. C., M. M. Schmitgen, C. Fuchs, C. Roth, R. C. Wolf, K. Bertsch, H. Flor, V. Grinevich and S. Boll (2019). “Oxytocin effects on pain perception and pain anticipation.” The Journal of Pain 20(10): 1187–1198.

Horta, M., D. Pehlivanoglu and N. C. Ebner (2020). “The role of intranasal oxytocin on social cognition: an integrative human lifespan approach.” Current behavioral neuroscience reports 7: 175–192.

Horta, M., R. Polk and N. C. Ebner (2023). “Single dose intranasal oxytocin administration: Data from healthy younger and older adults.” Data in Brief 51: 109669.

Horta, M., M. Ziaei, T. Lin, E. C. Porges, H. Fischer, D. Feifel, R. N. Spreng and N. C. Ebner (2019). “Oxytocin alters patterns of brain activity and amygdalar connectivity by age during dynamic facial emotion identification.” Neurobiology of aging 78: 42–51.

Hu, Y., D. Scheele, B. Becker, G. Voos, B. David, R. Hurlemann and B. Weber (2016). “The effect of oxytocin on third-party altruistic decisions in unfair situations: An fMRI study.” Scientific reports 6(1): 20236.

Ionta, S., L. Heydrich, B. Lenggenhager, M. Mouthon, E. Fornari, D. Chapuis, R. Gassert and O. Blanke (2011). “Multisensory mechanisms in temporo-parietal cortex support self-location and first-person perspective.” Neuron 70(2): 363–374.

Jenkinson, M., P. Bannister, M. Brady and S. Smith (2002). “Improved optimization for the robust and accurate linear registration and motion correction of brain images.” Neuroimage 17(2): 825–841.

Jenkinson, M. and S. Smith (2001). “A global optimisation method for robust affine registration of brain images.” Medical image analysis 5(2): 143–156.

Jiang, W., L. Cai and Z. Wang (2023). “Common hyper-entropy patterns identified in nicotine smoking, marijuana use, and alcohol use based on uni-drug dependence cohorts.” Medical & biological engineering & computing 61(12): 3159–3166.

Kawasaki, M., A. Sakai and Y. Ueta (2024). “Pain Modulation by Oxytocin.” Peptides: 171263.

Kheradmand, A. and A. Winnick (2017). “Perception of upright: multisensory convergence and the role of temporo-parietal cortex.” Frontiers in neurology 8: 552.

Krall, S. C., C. Rottschy, E. Oberwelland, D. Bzdok, P. T. Fox, S. B. Eickhoff, G. R. Fink and K. Konrad (2015). “The role of the right temporoparietal junction in attention and social interaction as revealed by ALE meta-analysis.” Brain Structure and Function 220: 587–604.

Lee, H.-J., A. H. Macbeth, J. H. Pagani and W. S. Young 3rd (2009). “Oxytocin: the great facilitator of life.” Progress in neurobiology 88(2): 127–151.

Leng, G. and M. Ludwig (2016). “Intranasal oxytocin: myths and delusions.” Biological psychiatry 79(3): 243–250.

Li, Z., Z. Fang, N. Hager, H. Rao and Z. Wang (2016). “Hyper-resting brain entropy within chronic smokers and its moderation by Sex.” Scientific reports 6(1): 29435.

Li, Z., Y. Zhu, A. R. Childress, J. A. Detre and Z. Wang (2012). “Relations between BOLD fMRI-derived resting brain activity and cerebral blood flow.”

Lin, C., S.-H. Lee, C.-M. Huang, G.-Y. Chen, P.-S. Ho, H.-L. Liu, Y.-L. Chen, T. M.-C. Lee and S.-C. Wu (2019). “Increased brain entropy of resting-state fMRI mediates the relationship between depression severity and mental health-related quality of life in late-life depressed elderly.” Journal of affective disorders 250: 270–277.

Lin, L., D. Chang, D. Song, Y. Li and Z. Wang (2022). “Lower resting brain entropy is associated with stronger task activation and deactivation.” NeuroImage 249: 118875.

Liu, P.-S., D.-H. Song, X.-P. Deng, Y.-Q. Shang, G. Qiu, Z. Wang and H. Zhang (2024). “Intermittent theta burst stimulation (iTBS)-induced changes of resting-state brain entropy (BEN).” bioRxiv: 2024.2005. 2015.591015.

Liu, P., T. Lin, D. Feifel and N. C. Ebner (2022). “Intranasal oxytocin modulates the salience network in aging.” NeuroImage 253: 119045.

Liu, X., D. Song, Y. Yin, C. Xie, H. Zhang, H. Zhang, Z. Zhang, Z. Wang and Y. Yuan (2020). “Altered Brain Entropy as a predictor of antidepressant response in major depressive disorder.” Journal of Affective Disorders 260: 716–721.

Lussier, D., Y. Cruz-Almeida and N. C. Ebner (2019). “Musculoskeletal pain and brain morphology: oxytocin’s potential as a treatment for chronic pain in aging.” Frontiers in Aging Neuroscience 11: 338.

Markiewicz, C. J., K. J. Gorgolewski, F. Feingold, R. Blair, Y. O. Halchenko, E. Miller, N. Hardcastle, J. Wexler, O. Esteban and M. Goncalves (2021). “OpenNeuro: An open resource for sharing of neuroimaging data.” BioRxiv: 2021.2006. 2028.450168.

Martins, D., K. Brodmann, M. Veronese, O. Dipasquale, N. Mazibuko, U. Schuschnig, F. Zelaya, A. Fotopoulou and Y. Paloyelis (2022). ““Less is more”: a dose-response account of intranasal oxytocin pharmacodynamics in the human brain.” Progress in neurobiology 211: 102239.

Neumann, I. D. (2008). “Brain oxytocin: a key regulator of emotional and social behaviours in both females and males.” Journal of neuroendocrinology 20(6): 858–865.

Power, J. D., K. A. Barnes, A. Z. Snyder, B. L. Schlaggar and S. E. Petersen (2012). “Spurious but systematic correlations in functional connectivity MRI networks arise from subject motion.” Neuroimage 59(3): 2142–2154.

Quintana, D., L. T. Westlye, S. Hope, T. Naerland, T. Elvsåshagen, E. Dørum, Ø. Rustan, M. Valstad, L. Rezvaya and H. Lishaugen (2017). “Dose-dependent social-cognitive effects of intranasal oxytocin delivered with novel Breath Powered device in adults with autism spectrum disorder: a randomized placebo-controlled double-blind crossover trial.” Translational psychiatry 7(5): e1136–e1136.

Quintana, D. S., A. Lischke, S. Grace, D. Scheele, Y. Ma and B. Becker (2021). “Advances in the field of intranasal oxytocin research: lessons learned and future directions for clinical research.” Molecular Psychiatry 26(1): 80–91.

Rash, J. A., A. Aguirre-Camacho and T. S. Campbell (2014). “Oxytocin and pain: a systematic review and synthesis of findings.” The Clinical journal of pain 30(5): 453–462.

Rey, A. (1958). “L’examen clinique en psychologie.”

Richman, J. S. and J. R. Moorman (2000). “Physiological time-series analysis using approximate entropy and sample entropy.” American journal of physiology-heart and circulatory physiology 278(6): H2039–H2049.

Rocchetti, M., J. Radua, Y. Paloyelis, L. A. Xenaki, M. Frascarelli, E. Caverzasi, P. Politi and P. FusarLPoli (2014). “Neurofunctional maps of the ‘maternal brain’and the effects of oxytocin: A multimodal voxelLbased metaLanalysis.” Psychiatry and clinical neurosciences 68(10): 733–751.

Shahrestani, S., A. H. Kemp and A. J. Guastella (2013). “The impact of a single administration of intranasal oxytocin on the recognition of basic emotions in humans: a meta-analysis.” Neuropsychopharmacology 38(10): 1929–1936.

Song, D.-H., X.-P. Deng, Y.-Q. Shang, D. Chang and Z. Wang (2024). “Altered resting-state brain entropy (BEN) by rTMS across the human cortex.” bioRxiv: 2024.2007. 2016.601109.

Song, D.-H. and Z. Wang (2024). “Regional Brain Entropy During Movie-watching.” bioRxiv: 2024.2006. 2012.598767.

Song, D.-H. and Z. Wang (2024). “The Relationships of Resting-state Brain Entropy (BEN), Ovarian Hormones and Behavioral Inhibition and Activation Systems (BIS/BAS).” bioRxiv: 2024.2006. 2004.595915.

Song, D., D. Chang, J. Zhang, Q. Ge, Y.-F. Zang and Z. Wang (2019). “Associations of brain entropy (BEN) to cerebral blood flow and fractional amplitude of low-frequency fluctuations in the resting brain.” Brain imaging and behavior 13(5): 1486–1495.

Song, D., D. Chang, J. Zhang, W. Peng, Y. Shang, X. Gao and Z. Wang (2019). “Reduced brain entropy by repetitive transcranial magnetic stimulation on the left dorsolateral prefrontal cortex in healthy young adults.” Brain imaging and behavior 13: 421–429.

Song, D., K. Jann and D. J. Wang (2024). “Methodological development and applications of nonlinear dynamic analysis for neuroimaging.” Frontiers in Neurology 15: 1387356.

Stam, C. J. (2005). “Nonlinear dynamical analysis of EEG and MEG: review of an emerging field.” Clinical neurophysiology 116(10): 2266–2301.

Tracy, L. M., N. Georgiou-Karistianis, S. J. Gibson and M. J. Giummarra (2015). “Oxytocin and the modulation of pain experience: Implications for chronic pain management.” Neuroscience & Biobehavioral Reviews 55: 53–67.

Valstad, M., G. A. Alvares, M. Egknud, A. M. Matziorinis, O. A. Andreassen, L. T. Westlye and D. S. Quintana (2017). “The correlation between central and peripheral oxytocin concentrations: a systematic review and meta-analysis.” Neuroscience & Biobehavioral Reviews 78: 117–124.

Van Overwalle, F. (2009). “Social cognition and the brain: a metaLanalysis.” Human brain mapping 30(3): 829–858.

Wang, Y., A. Metoki, Y. Xia, Y. Zang, Y. He and I. R. Olson (2021). “A large-scale structural and functional connectome of social mentalizing.” NeuroImage 236: 118115.

Wang, Z. (2021). “The neurocognitive correlates of brain entropy estimated by resting state fMRI.” NeuroImage 232: 117893.

Wang, Z. and A. s. D. N. Initiative (2020). “Brain entropy mapping in healthy aging and Alzheimer’s disease.” Frontiers in Aging Neuroscience 12: 596122.

Wang, Z., Y. Li, A. R. Childress and J. A. Detre (2014). “Brain entropy mapping using fMRI.” PloS one 9(3): e89948.

Wechsler, D. (1981). “WAIS-R: Manual: Wechsler adult intelligence scale-revised.” (No Title).

Xue, S.-W., Q. Yu, Y. Guo, D. Song and Z. Wang (2019). “Resting-state brain entropy in schizophrenia.” Comprehensive Psychiatry 89: 16–21.

Young, L., D. Dodell-Feder and R. Saxe (2010). “What gets the attention of the temporo-parietal junction? An fMRI investigation of attention and theory of mind.” Neuropsychologia 48(9): 2658–2664.

Zhang, Y., M. Brady and S. Smith (2001). “Segmentation of brain MR images through a hidden Markov random field model and the expectation-maximization algorithm.” IEEE transactions on medical imaging 20(1): 45–57.

Zhao, Z., Y. Shuai, Y. Wu, X. Xu, M. Li and D. Wu (2024). “Age-dependent functional development pattern in neonatal brain: an fMRI-based brain entropy study.” NeuroImage: 120669.

Zou, Q.-H., C.-Z. Zhu, Y. Yang, X.-N. Zuo, X.-Y. Long, Q.-J. Cao, Y.-F. Wang and Y.-F. Zang (2008). “An improved approach to detection of amplitude of low-frequency fluctuation (ALFF) for resting-state fMRI: fractional ALFF.” Journal of neuroscience methods 172(1): 137–141.

Zunhammer, M., S. Geis, V. Busch, M. W. Greenlee and P. Eichhammer (2015). “Effects of intranasal oxytocin on thermal pain in healthy men: a randomized functional magnetic resonance imaging study.” Psychosomatic medicine 77(2): 156–166.

